# Pulses of Class I PI3kinase activity identify the release and recapture of prey from neutrophil phagosomes

**DOI:** 10.1101/2024.02.29.582694

**Authors:** Clare F Muir, Constantino Carlos Reyes-Aldasoro, Felix E Ellett, Tomasz K Prajsnar, Yin X Ho, Audrey Bernut, Catherine A Loynes, Stone Elworthy, Kieran A Bowden, Ashley J Cadbury, Lynne R Prince, Jason S King, Alison M Condliffe, Steven A Renshaw

## Abstract

Class I PI3kinases coordinate the delivery of microbicidal effectors to the phagosome by forming the phosphoinositide lipid second messenger, phosphatidylinositol (3, 4, 5)-trisphosphate (PIP3). However, the dynamics of PIP3 in neutrophils during a bacterial infection are unknown. We have therefore developed an *in vivo,* live zebrafish infection model that enables visualisation of dynamic changes in Class 1 PI3kinases (PI3K) signalling on neutrophil phagosomes in real-time. We have identified that on approximately 12% of neutrophil phagosomes PHAkt-eGFP, a reporter for Class 1 PI3K signalling, re-recruits in pulsatile bursts. This phenomenon occurred on phagosomes containing structurally and morphologically distinct prey, including *Staphylococcus aureus* and *Mycobacterium abscessus*, and was dependent on the activity of the Class 1 PI3K isoform, PI3kinase γ. Detailed imaging suggested that ‘pulsing phagosomes’ represent neutrophils transiently reopening and reclosing phagosomes. This finding challenges the concept that phagosomes remain closed after prey engulfment and we propose that neutrophils occasionally use this alternative pathway of phagosome maturation to release phagosome contents and/or to restart phagosome maturation if digestion has stalled.

## Introduction

Neutrophils are the most abundant white blood cells in the human body and are crucial in the immune system’s early defence against bacterial infection. Neutrophils are avid phagocytes, capturing bacteria in membrane-bound vesicles called phagosomes for subsequent execution and digestion. The production of phosphatidylinositol (3, 4, 5)-trisphosphate (PIP3) on phagosome membranes by Class 1 PI3kinases (PI3K) facilitates intra-phagosomal killing of bacteria by co-ordinating reactive oxygen species (ROS) formation ^1,2^, degranulation ^3,4^ and actin dynamics, enabling the phagosome to form and fuse with other organelles ^5,6^. However, although PIP3 has been shown to enable phagosome formation around large, inert prey such as complement component 3 fragment (C3bi)-opsonised zymosan and IgG-opsonised erythrocytes in neutrophil and macrophage-like cells ^7,8^ it is unknown how regulation of PIP3 in space and time controls the digestion of bacterial-containing phagosomes. It is also unknown which PI3K isoform generates PIP3 on neutrophil phagosomes. The Pleckstrin Homology domain of Akt (PHAkt) recognises PIP3 and also PI(3,4)P2 (the degradation product of PIP3) and has been used extensively to characterise Class 1 PI3K activity during phagocytosis by macrophage-like cell lines ^9^. By developing an *in vivo* infection model using the transgenic zebrafish neutrophil reporter *Tg(lyz:PHAkt-eGFP)i277* ^10^, we show that PIP3/PI(3,4)P2 persists on neutrophil phagosomes far longer than *in vitro* macrophage models report and that some PIP3/PI(3,4)P2 positive phagosomes repeatedly reopen and reclose, associated with re-recruitment of the reporter in a pulsatile manner. ‘Pulsing’ occurs when both live and dead *Staphylococcus aureus* are ingested, as well as *Mycobacterium abscessus* and inert beads, and is dependent on PI3kinase γ. We suggest that ‘pulsing phagosomes’ are indicative of repeated cycles of neutrophils releasing and recapturing prey. This process may offer neutrophils an opportunity to restart phagosome maturation, enhancing pathogen digestion, and also to potentially modulate local inflammation by communicating to neighbouring cells what has been eaten ^11^.

## Results

### 1. PHAkt-eGFP recruits in pulsatile bursts to neutrophil phagosomes

To investigate the role of PIP3/PI(3,4)P2 during phagocytosis, we injected *Staphylococcus aureus,* an important pathogen of humans and animals, into the zebrafish neutrophil PIP3/PI(3,4)P2 reporter line, *Tg(lyz:PHAkt-eGFP)i277* ^12^ (Fig. 1A). Minimal PHAkt-eGFP recruited during early formation of the phagocytic cup. PHAkt-eGFP recruited uniformly to the cup prior to closure and then localised at sites of cup closure (Movie S1). PHAkt-eGFP then gradually diminished on the majority of phagosomes on average over 17.9mins (+/− SD 17.9) (Fig. 1B-E). However, we observed to our surprise that in a subset of phagosomes, PHAkt-eGFP then re-recruited to the phagosome membrane in pulsatile bursts (Movie S2). A ‘pulse’ was defined as a transient but intense surge in PHAkt-eGFP recruitment to the phagosome (Fig.1F-G, Movie S2). Repeated pulsing occurred on 12.4% (+/− SD 24.9) of phagosomes within a neutrophil and 35.8% (+/− SD 36.7) of all neutrophils had at least one pulsing phagosome. Cycles of PHAkt-eGFP re-recruitment often occurred multiple times (Fig. 1H), typically x4 on each phagosome, but with up to 21 pulses observed on a phagosome containing a single bacteria which had been ‘shuttled’ ^13^ from another neutrophil. Some neutrophils phagocytosed more bacteria than others and we wondered whether these ‘hungry phagocytes’ had more pulsing phagosomes, perhaps reflecting exhaustion of phagocytic capacity ^14^. However, there was no correlation between the number of pulses and the number of bacteria within a neutrophil for live bacteria (Spearman’s rank correlation= −0.3091, p=0.0907) or dead bacteria (Spearman’s rank correlation=0.1447. p=0.3998) (Fig.1I).

**Fig. 1.**
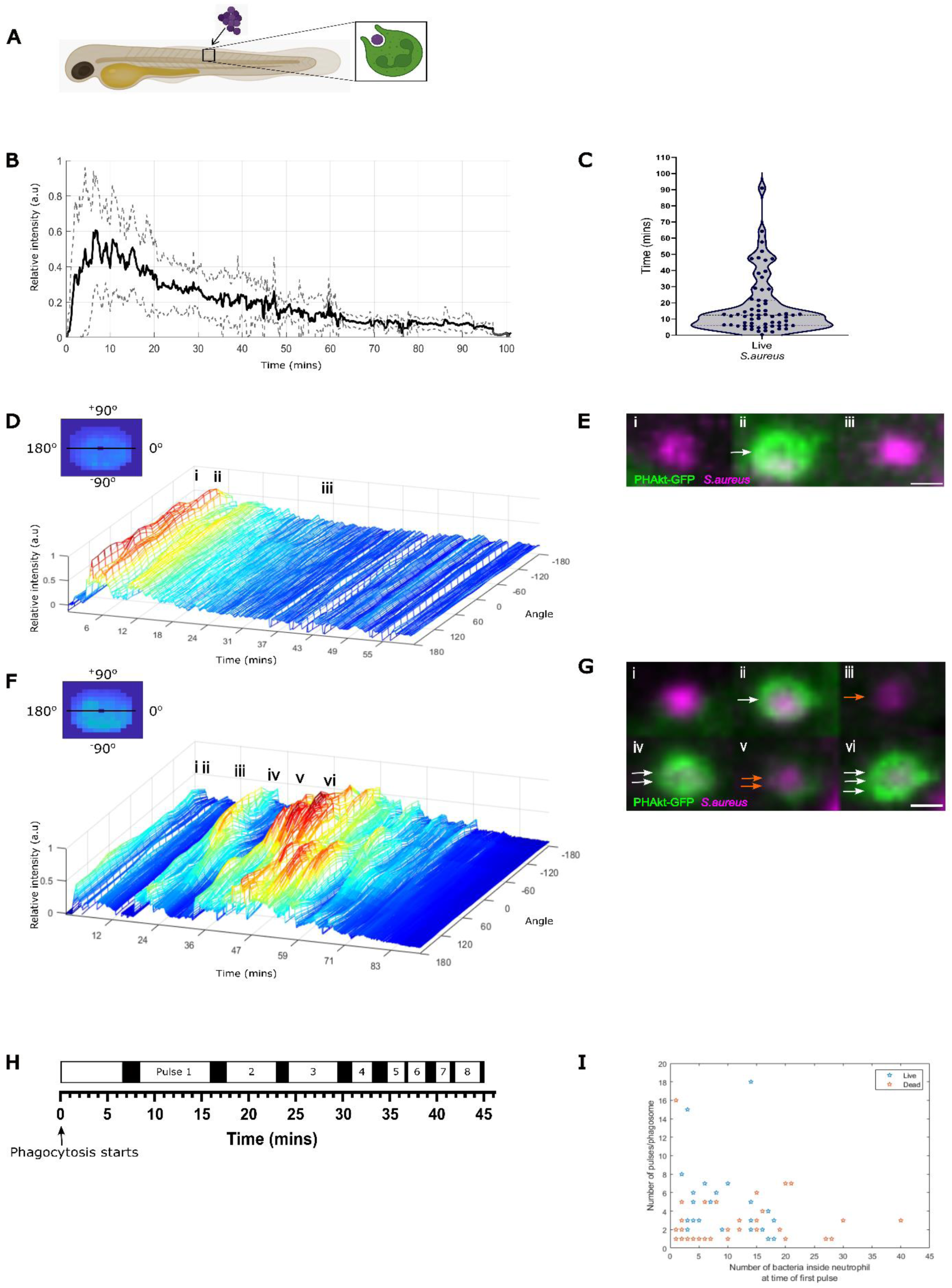
PHAkt-eGFP recruits in pulsatile bursts to neutrophil phagosomes. **(A)** Schematic illustrating location of *S.aureus* injection into day 3 *Tg(lyz:PHAkt-EGFP)i277* zebrafish larvae. **(B)** Quantification of PHAkt-eGFP fluorescence on the phagosome membrane during phagocytosis of pHrodo™ Red *S.aureus*. Data shown is the average intensity values of 9 phagosomes from 4 experiments +/− SD. **(C)** Violin plot showing the duration of PHAkt-eGFP recruitment to neutrophil phagosomes following phagocytosis of live *S.aureus*. Data shown is the Median with the 25th and 75th percentiles (63 phagosomes analysed, 11 experiments). **(D)** Quantification of PHAkt-eGFP fluorescence to the phagosome membrane over time. **(E)** Sequential images of a neutrophil phagosome. i-iii correlate to Fig.1D. i. Start of phagocytosis. ii. Surge of PHAkt-eGFP recruitment as the phagosome closes (white arrow). iii. PHAkt-eGFP diminishes from phagosome. Scale Bar = 1µm. **(F)** Quantification of PHAkt-eGFP fluorescence on a pulsing phagosome. **(G)** Sequential images of a pulsing phagosome. i-vi correlate to Fig.1G. i. Start of phagocytosis. ii. Surge of PHAkt-eGFP recruitment as the phagosome closes (white arrow). iii. PHAkt-eGFP diminishes from phagosome (orange arrow). iv. 1^st^ pulse (two white arrows). v. Loss of PHAkt-eGFP recruitment (two orange arrows). vi 2^nd^ pulse (three white arrows). Scale Bar = 1µm. **(H)** Schematic illustrating pulsatile recruitment of PHAkt-eGFP to phagosomes. Data represent average values from multiple phagosomes, 11 experiments. **(I)** Pulses occur irrespective of the number of bacteria within a neutrophil. Live *S.aureus*: 31 phagosomes analysed. Spearman’s rank correlation between the number of bacteria inside neutrophil at time of 1st pulse and number of pulses/phagosome was - 0.3091 with a corresponding p-value of 0.0907. Dead *S.aureus*: 36 phagosomes analysed. Spearman’s rank correlation between the number of bacteria inside neutrophil at time of 1st pulse and number of pulses/phagosome was 0.1447 with a corresponding p-value of 0.3998.

### 2. Pulsatile bursts of PHAkt-eGFP recruitment are a neutrophil response to prey

Having observed that PHAkt-eGFP re-recruits to phagosomes containing live *S.aureus*, we wondered if this phenomenon reflected bacterial manipulation of PIP3/PI(3,4)P2 signalling. To investigate this, *Tg(lyz:PHAkt-eGFP)i277* larvae were injected with heat-killed *S.aureus* (Fig. 2A), hypothesising that pulsing would not be observed if it was due to an active bacterial response to neutrophil attack. However, no significant difference was identified between the percentage of pulsing phagosomes in neutrophils containing live (12.4% +/− SD 24.9) vs dead bacteria (9.3% +/− SD 19.0). Pulsing therefore does not reflect active manipulation of PIP3/PI(3,4)P2 signalling by *S.aureus*.

**Fig. 2.**
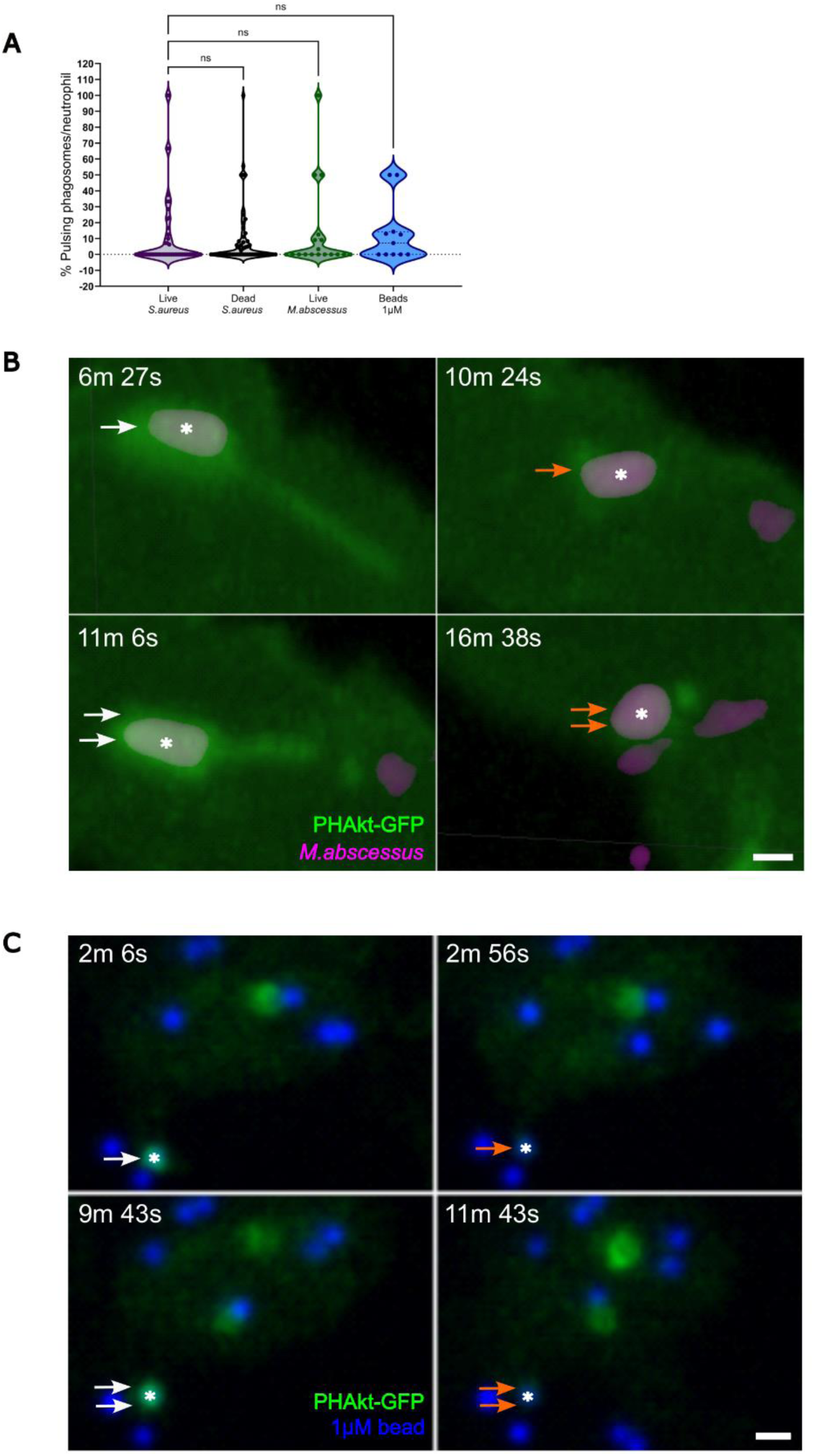
Pulsatile bursts of PHAkt-eGFP recruitment are a neutrophil response to prey. **(A)** Violin plot showing the % of ‘pulsing’ phagosomes for live and dead *S.aureus*, live *M.abscessus* and 1µM beads. Data shown is the Median with the 25^th^ and 75^th^ percentiles. Live *S.aureus* = 46 phagosomes, 11 experiments. Dead *S.aureus* = 54 phagosomes, 10 experiments. Live *M.abscessus* =18 phagosomes, 6 experiments. 1µM beads= 11 phagosomes, 4 experiments. **(B)** ‘Pulses’ occur on phagosomes containing *M.abscessus*. PHAkt-eGFP recruits to phagosomes during phagocytosis (white arrow) before gradually diminishing (orange arrow). PHAkt-eGFP then re-recruits to the phagosome (‘1st pulse’) (two white arrows) and then diminishes again (two orange arrows). Scale Bar = 2µm. **(C)** ‘Pulses’ occur on phagosomes containing 1µM beads. PHAkt-eGFP recruits to phagosomes during phagocytosis (white arrow) before gradually diminishing (orange arrow). PHAkt-eGFP then re-recruits to the phagosome (‘1st pulse’) (two white arrows) and then diminishes again (two orange arrows). Phagocytosis starts at 0mins. Scale Bar = 2µm.

To investigate whether phagosomal ‘pulsing’ is a specific response to *S.aureus*, larvae were injected with a structurally and morphologically distinct prey, *Mycobacterium abscessus* (Fig.2A and B, Movie S3). Pulsatile recruitment of PHAkt-eGFP was again identified and the frequency of pulses was similar to phagosomes containing *S.aureus* (13% +/− SD 26.9) (Fig. 2A), indicating that PHAkt-eGFP re-recruitment occurs irrespective of the species of bacteria engulfed. To distinguish if pulsing is a neutrophil response to phagocytosis of bacteria *vs* non-bacterial prey, larvae were injected with 1µm polystyrene beads (F13083, a size similar to *S.aureus* (Fig.2C: Movie S4). Again, a similar frequency of pulsing was identified (13.36% (+/− SD 19.02) (Fig. 2A). Together these data suggest that ‘pulsing’ is a normal neutrophil response to prey, rather than bacteria attempting to manipulate host signalling.

### 3. Pulsatile recruitment of PHAkt-eGFP is associated with incomplete phagocytosis, bacterial escape and pathogen shuttling

During our experiments, we observed some neutrophils expelling *S.aureus* from PHAkt-eGFP positive phagosomes (Movie S5). Neutrophils often re-phagocytosed the same bacteria, and in this setting PHAkt-eGFP once again recruited to the enveloping membranes. We therefore considered whether pulsing phagosomes reflected phagosomes reopening, releasing phagosome contents and then recapturing. Imaging of this phenomenon was challenging as phagosomes often only opened transiently and via a small pore. However, 3D reconstruction of ‘pulsing phagosomes’ (Fig.3A and Movie S6) enabled us to confirm that at least 87.1% (27/31) of ‘pulsing phagosomes’ did indeed reopen. On reopening, PHAkt-eGFP recruitment rapidly dropped, with the bacteria usually remaining within the phagocytic cup, before the phagosome resealed (Fig.3A and Movie S6). In 5.1% (7/138) ‘pulses’, bacteria were fully released from the phagocytic cup but remained close to the neutrophil surface before being recaptured. 94% (29/31) of bacteria from ‘pulsing’ phagosomes were successfully recaptured by the same neutrophil. 3% (1/31) bacteria were released and then re-phagocytosed by a non-fluorescent phagocytic cell (presumed macrophage) and 3% (1/31) bacteria was shuttled into another neutrophil. To confirm that pulsing phagosomes re-open, actin and PHAkt-eGFP dynamics were assessed in parallel using larvae from a cross of *Tg(lyz:PHAkt-eGFP)i277* and *Tg(mpx:Lifeact-Ruby)sh608* (Fig.3B). This confirmed that PHAkt-eGFP dissociates from the phagosome when phagosomes re-open and release bacteria and that PHAkt-eGFP re-recruits to phagosomes as a ‘pulse’ when the same bacteria are recaptured. We therefore propose that ‘pulses’ indicate phagosomes which reopen and then recapture prey.

**Fig. 3.**
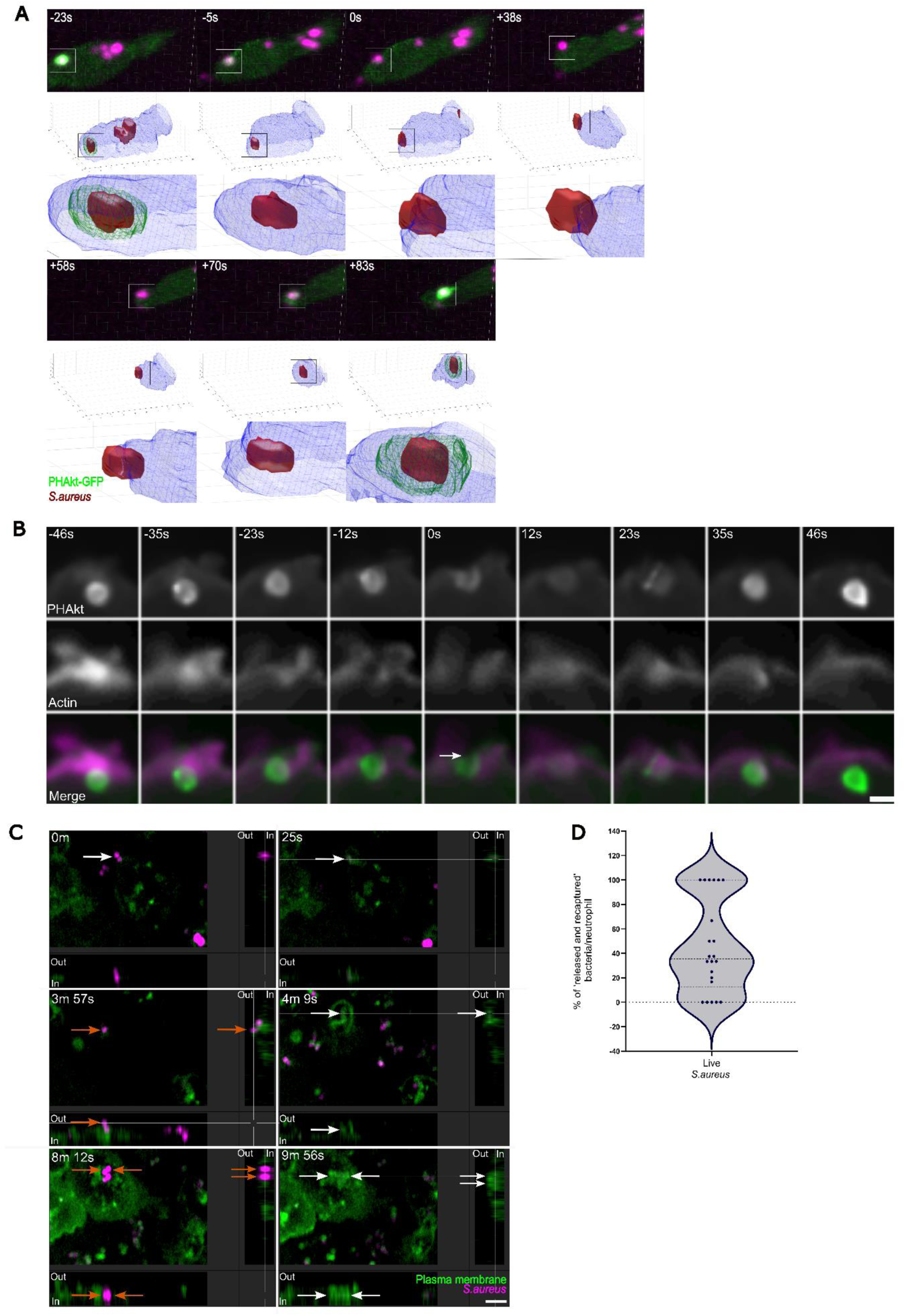
Pulses of PHAkt-eGFP are repeated episodes of bacteria being ‘released and recaptured’ from phagosomes. **(A)** 3D reconstruction of PHAkt-eGFP dynamics when *S.aureus* is released and recaptured from a phagosome. 0s is the time that the bacteria is rephagocytosed. **(B)** Sequential images capturing the dynamics of PHAkt-eGFP and actin when bacteria are released and recaptured from a phagosome. Arrow shows separation of cortical actin filaments as pulsing phagosome reopens onto the neutrophil surface. Scale Bar = 2µm. **(C)** Orthogonal views from a timelapse illustrating that human neutrophils release and recapture *S.aureus* from phagosomes. 0mins = start of phagocytosis. *S.aureus* is engulfed into a phagosome (25s). *S.aureus* is released from a phagosome into the extracellular space (orange arrows). *S.aureus* is then recaptured (white arrows) (1^st^ pulse). The phagosome then re-opens (2 orange arrows) and then recloses (2 white arrows) (2^nd^ pulse). Scale Bar = 5µm. **(D)** Violin plot showing the % of ‘release and recapture’ events in human neutrophils. Data shown is the Median with the 25th and 75th percentiles. 140 phagosomes analysed, 2 experiments.

### 4. Human neutrophils release and recapture prey

Having identified that some pulsing phagosomes reopen, we wished to identify whether this phenomenon also occurred in human neutrophils, and to exclude the possibility that the PHAkt-eGFP construct might exert a dominant negative effect on normal PIP3/PI(3,4)P2 signalling and thereby hinder effective phagocytosis. To address this, we stained the membrane of primary human neutrophils with CellMask™ Deep Red and imaged them phagocytosing pHrodo™ Green stained *S.aureus* (Movie S7). This demonstrated that 45.6% (+/− SD 38.5) of human neutrophil phagosomes also release and recapture bacteria (Fig.3C and D).

### 5. Pulses of PHAkt-eGFP are abolished by PI3K γ inhibition

During imaging, we identified that PHAkt-eGFP uniformly increases on the phagosome membrane as the phagocytic cup closes and that there is a second increase in PHAkt-eGFP recruitment at the neck of the phagosome, prior to sealing. We next aimed to identify the Class 1 PI3Kinase enzyme which drives PIP3 production on neutrophil phagosomes ^15^. The three tyrosine kinase-linked Class1A PI3kinase isoforms are α, β, and δ, and the sole G-protein activated Class1B PI3kinase isoform is PI3K γ ^15^. Neutrophils express abundant PI3kinases γ and δ ^15^ and, in zebrafish, inhibition of PI3kinase γ completely abolishes the migration of neutrophils to an injured tail fin ^16^. Although unexplored in a zebrafish infection model, PI3kinase δ inhibition has been reported to reduce the migration of human neutrophils *in vitro* ^17^ and also reduces neutrophil migration towards a TNFα injection site in a mouse ^18^. This makes it difficult to assess whether neutrophils from transgenic fish lacking PI3kinase γ or δ phagocytose prey *in vivo*, as such neutrophils will not migrate to the infection site. To assess the role of PI3kinases γ and δ in phagocytosis, neutrophils were therefore allowed to migrate to the infection site for 60mins and then zebrafish were exposed to class 1 PI3kinase inhibitors. Based on our published previous work, we incubated larvae for 30mins with 100μM of the PI3kinase δ inhibitor, CAL-101 ^19^) before immediately commencing imaging. In both control zebrafish and zebrafish exposed to CAL-101, pulsatile bursts of PHAkt-eGFP recruitment occurred on neutrophil phagosomes (Fig.4A-C).

**Fig. 4.**
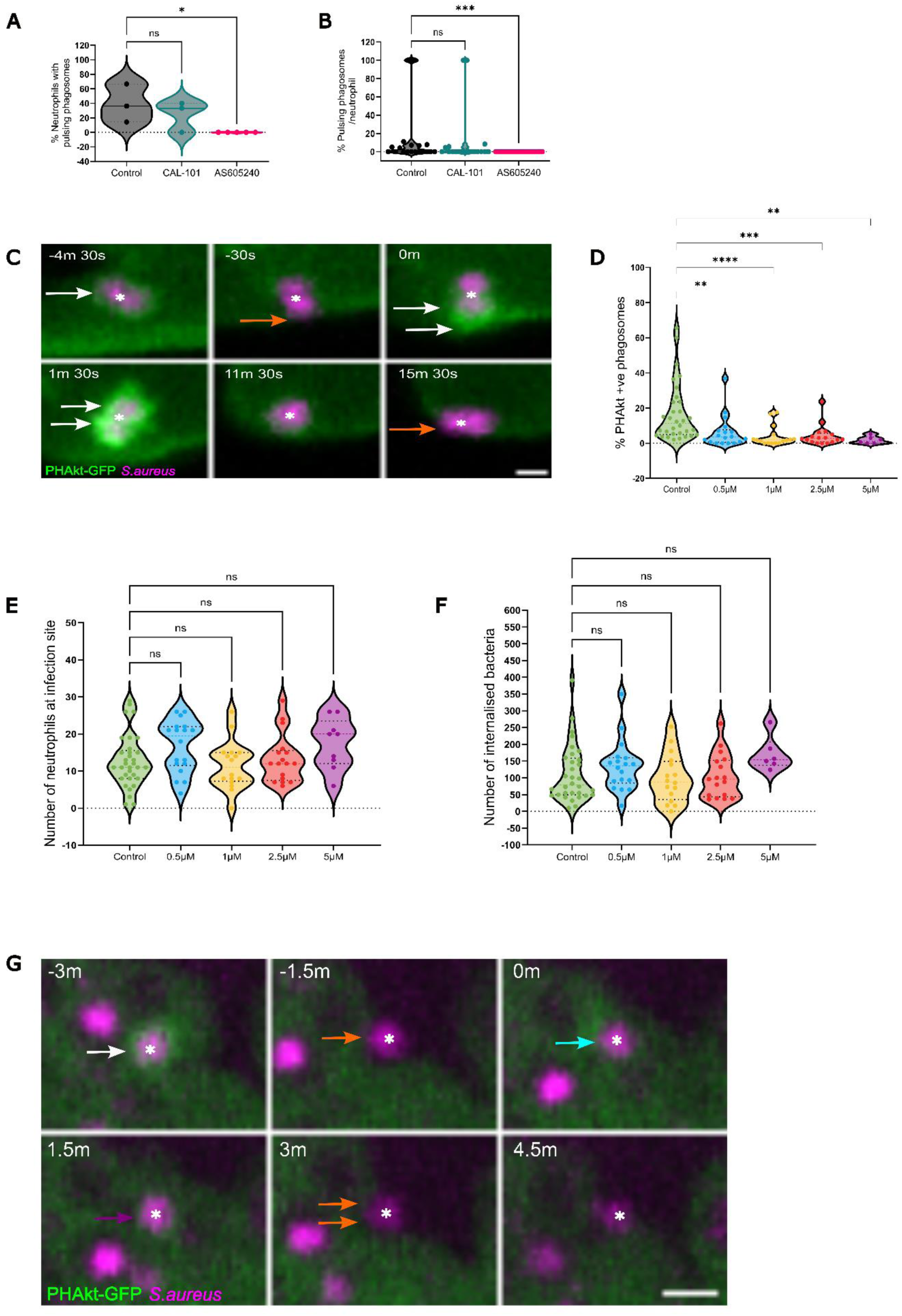
Pulses of PHAkt occur despite inhibition with CAL-101 and are abolished by AS605240. **(A)** Violin plot showing the % of neutrophils that have pulsing phagosomes. Timelapse started 2hours post infection following 30min incubation of *Tg(lyz:PHAkt-EGFP)i277* larvae with 100µM CAL-101 and 1µM AS605240. Data shown is the Median with the 25th and 75th percentiles from 87 neutrophils, 11 experiments. **(B)** Violin plot showing % of pulsing phagosomes in each neutrophil in a timelapse starting 2hours post infection following 30min incubation of *Tg(lyz:PHAkt-eGFP)i277* larvae with 100µM CAL-101 and 1µM AS605240. Data shown is the Median with the 25th and 75th percentiles from 780 phagosomes, 11 experiments. **(C)** Sequential images illustrating pulsatile recruitment of PHAkt-eGFP to a phagosome exposed to 100µM CAL-101. Scale Bar = 2µm. **(D)** Violin plot showing quantification of the % of PHAkt-eGFP +ve phagosomes 2hours post infection following 30mins incubation with AS605240. Data shown is the median with the 25th and 75th percentiles from 6 experiments. **(E)** Violin plot showing quantification of the number of neutrophils at a *S.aureus* infection site 2hours post infection following 30mins incubation with AS605240. Data shown is the median with the 25th and 75th percentiles from 6 experiments. **(F)** Violin plot showing quantification of the number of bacteria/neutrophil 2hours post infection following 30mins incubation with AS605240. Data shown is the median with the 25th and 75th percentiles from 6 experiments. **(G)** Sequential images illustrating that PHAkt-eGFP recruits to neutrophil phagosomes prior to exposure to 1µM AS605240 (white arrow). Bacteria is then released from the phagosome (orange arrow). Following exposure to 1µM AS605240, the neutrophil attempts to rephagocytose the bacteria (blue arrow) but PHAkt-eGFP does not recruit to the phagosome (magenta arrow) and the bacteria is released from the phagosome (two orange arrows). Scale Bar = 2µm.

We titrated the dose of PI3K γ inhibitor AS605240 to identify that a delayed 30mins incubation with 1μM of this compound significantly decreased the % of PHAkt-eGFP positive phagosomes, but not the number of neutrophils recruited to the infection site nor the number of bacteria internalised per neutrophil (Fig.4D-F). This suggested to us that neutrophils recruited to the infection site and phagocytosed bacteria in the 60mins before the PI3K γ inhibitor was applied. However, the subsequent application of AS605240 then reduced the production of PIP3/PI(3,4)P2 on neutrophil phagosomes (Fig.4G: Movie S8). Close examination of time-lapses of AS605240-treated larvae showed that although some PHAkt-eGFP recruited to neutrophils as they initiated phagocytosis, the later larger surges of PHAkt-eGFP recruitment, firstly to the fully formed phagocytic cup and secondly, to the neck of the phagosome during closure, did not occur. In the absence of PHAkt-eGFP recruitment to phagosomes, the partially phagocytosed bacteria were released back into the tissue. Occasionally, neutrophils attempted to re-phagocytose the bacteria but again, visible re-recruitment of PHAkt-eGFP (‘pulsing’) was absent (Fig.4A, B and G: Movie S8). We, therefore, concluded that PI3kinase γ enables the majority of PHAkt-eGFP recruitment to neutrophil phagosomes. We propose that without functional PI3kinase γ activity, neutrophil phagosomes fail to close effectively, and prey is subsequently released into the tissue but not re-captured.

## Discussion

Using an *in vivo* model of Class 1 PI3K signalling, we have identified an intriguing new phenomenon by which neutrophils repeatedly release and recapture prey. This phenomenon occurs following phagocytosis of structurally and morphologically distinct bacteria as well as inert beads, and it is observed in human neutrophils.

In the overwhelming majority of phagosomes, we were unable to visualise continuity between the phagosome and cell membrane and it is therefore plausible that pulsing phagosomes may have fully sealed and then re-fused with the plasma membrane to release prey (non-lytic exocytosis) ^20–22^.

Our findings differ from current understanding of non-lytic exocytosis which, in macrophage-like cells, does not occur with inert latex beads and occurs with less frequency with avirulent pathogens ^23,24^. Additionally, prey that has been expelled from neutrophils and macrophages by non-lytic exocytosis is permanently ejected, whereas we observed that the majority of prey released from pulsing phagosomes is recaptured- and often follows multiple release and recapture cycles ^23,24^. Non-lytic exocytosis is also reported to start 46mins-hours after phagocytosis ^23,24^, whereas we observed phagosome re-opening 10.5mins (+/− 12.49) after the start of phagocytosis.

Rather than permanently expelling a pathogen, pulsing phagosomes may instead enable neutrophils to release partially digested phagosome contents and this process may enable neutrophils to modulate the local inflammatory response. Greene *et al.* 2022 identified that phagolysosomes within primary human and murine macrophages and murine neutrophils repeatedly fused with the cell membrane and released soluble matter to the extracellular space. This process, termed ‘eructophagy’, enabled the activation of neighbouring cells that in turn amplifies local inflammation. Eructophagy was seen in approximately 10% of phagolysosomes over a 3-hour period, and was stimulated by inflammation and nutrient depletion. Eructophagy also decreased in bone marrow-derived macrophages from PI3kr6^−/−^ mice ^11^, lacking the gene for one of the regulatory subunits of PI3K γ, p84. The frequency of eructophagy and the conditions under which it occurs are similar to that which we describe for pulsing phagosomes, although eructophagy, to our knowledge, has not previously been described in human or zebrafish neutrophils.

Additionally, the membrane dynamics during eructophagy events have not been visualised ^11^ and here we show that some phagosomes re-open through a small pore whereas other phagosomes reopen entirely, releasing the bacteria onto the neutrophil surface before recapturing. Imaging quality was insufficient to distinguish whether 4/31 pulsing phagosomes reopened, but reopening was definitively identified for the remaining 27/31 events. Nevertheless, all pulsing phagosomes moved towards the periphery of the neutrophil before PHAkt-eGFP fluorescence diminished, as would be expected for a phagosome releasing contents into the tissue. Additionally, we identified that although 17% of neutrophil phagosomes fuse to form larger phagosomes, pulsing phagosomes never fused with other phagosomes. We therefore speculate that, as described for erutophagy ^11^, pulsing phagosomes preferentially fuse with the plasma membrane rather than to other phagosomes or, potentially, to other organelles. Alternatively, the failure of phagosomes to fuse with other organelles may represent a failure of maturation and potentially trigger reopening, similar to non-lytic exocytosis in which alkalinisation promotes phagosome reopening ^25,26^. It is also conceivable that the release and recapture of prey from pulsing phagosomes occurred because the phagosomes never sealed. Cech and Lehrer., 1984 described that 40% of neutrophil phagosomes containing *Candida albicans* remain open through ‘cleft-like micro channels’ ^27^. However, pulsing phagosomes normally pulse multiple times and we would not expect neutrophils to repeatedly fail to close a phagosome.

We identified that recruitment of PHAkt to neutrophil phagosomes is dependent on PI3kinase γ activity and that without PI3kinase γ neutrophils are unable to effectively retain prey. This suggests that PI3kinase γ, in addition to playing a key role in enabling neutrophil migration ^28^, might enable bacteria-containing phagosomes to seal. Supporting our findings, Quinn *et al.,* 2021 identified that Class 1PI3 kinases, which include PI3kinase γ, enable macropinosomes to seal ^29^ and potentially similar mechanisms may underpin phagocytosis of bacteria. There are two regulatory subunits for PI3kinase γ, namely p84 and p101, each of which co-ordinate different neutrophil responses ^30,31^. Future work may determine which PI3kinase γ regulator(s) enable PIP3 production on bacterial-containing phagosomes to facilitate closure. This is important as it may allow selective therapeutic targeting of neutrophil function. For example, although there are currently no p84/p101 inhibitors, selective nanobodies have recently been developed that can selectively target the different PI3kinase γ inhibitory complexes ^32^. Potentially these nanobodies could be used to prevent release and recapture events, thereby limiting local inflammation caused by the release of phagosomal contents into the extracellular matrix but without impacting neutrophil migration.

Our findings contrast with studies showing that macrophage-like cells do not require PI3kinases to phagocytose ‘bacterial-sized particles’ (<3μm IgG opsonized latex beads) ^5,33,35^. This may reflect differences in how PI3kinases coordinate phagocytosis in primary neutrophils *in vivo* vs macrophage-like cells *in vitro*. Additionally, activation of PI3kinase γ would be limited in studies using IgG opsonized latex beads because PI3kinase γ is principally activated by G protein-coupled receptors (e.g. complement receptors) not immunoglobulin receptors ^34,36^. Zebrafish larval neutrophils are perhaps more likely to use innate (e.g. complement), rather than adaptive, immune receptors to phagocytose prey, because the adaptive immune system is not fully developed until zebrafish are at least 4 weeks old ^38^. Our findings therefore suggest that there might be profound differences in how adaptive vs innate immune receptors co-ordinate phagocytosis and how neutrophils vs macrophages use class 1 PI3 kinases to phagocytose prey.

In conclusion, we have developed an *in vivo* model to image Class1 PI3K signalling in real-time during a bacterial infection. This model has enabled us to identify that PI3kinase γ produces the majority of PIP3/PI(3,4)P2 on bacterial phagosomes and that the majority of PI3kinase γ activity occurs after bacteria are internalised by the neutrophil. Additionally, pulses of class 1 PI3K activity occur when neutrophils repeatedly release and recapture prey from phagosomes, and this intriguing finding provides new insight into how neutrophils respond to bacteria during a live tissue infection.

## Supplementary Figures

**Figure S1 Dynamics of PHAkt-eGFP on neutrophil phagosomes following infection with *S.aureus*.** Minimal PHAkt-eGFP recruits during early formation of the phagocytic cup. PHAkt-eGFP recruits strongly at sites of cup closure (white arrows) and uniformly to the phagosome membrane after cup formation (white arrows). Green = PHAkt-eGFP. Magenta = pHrodo™ Red stained *S.aureus*.

**Figure S2 Pulsatile recruitment of PHAkt-eGFP to a neutrophil phagosome containing *S.aureus.*** Green = PHAkt-eGFP. Magenta = pHrodo™ Red stained S.aureus.

**Figure S3 Pulsatile recruitment of PHAkt-eGFP to a neutrophil phagosome containing *Mycobacterium abscessus*.** Green = PHAkt-eGFP. Magenta = pHrodo™ Red stained *M.abscessus*.

**Figure S4 Pulsatile recruitment of PHAkt-eGFP to a neutrophil phagosome containing 1µm latex beads (blue spheres).** 1µm bead is extracellular (in the tissue) at the start of the movie (white circle). A neutrophil phagocytoses the bead (1m 46s) and PHAkt-eGFP recruits to the phagosome membrane. The bead is then released from the phagosome (5m 7s). Neutrophil then rephagocytose *S.aureus* (5m 39s) (red dot) (1^st^ pulse). *S.aureus* is then released from the phagosome (7m 3s). Neutrophil then rephagocytose *S.aureus* (7m 26s) (red dot) (2^nd^ pulse).1µm latex beads (blue spheres).

**Figure S5 Neutrophil expelling and recapturing *S.aureus*.** A neutrophil phagocytoses *S.aureus* (12s) and PHAkt-eGFP recruits to the phagosome membrane. PHAkt-eGFP diminishes from the phagosome membrane and *S.aureus* is expelled from the phagosome onto the surface of the neutrophil (29s). *S.aureus* is recaptured by the neutrophil and PHAkt-eGFP re-recruits to the phagosome membrane (1st pulse) (33s). Green = PHAkt-eGFP. Magenta = pHrodo™ red stained *S.aureus*.

**Figure S6 3D reconstruction of a pulsing phagosome showing that *S.aureus* is repeatedly released and recaptured by the neutrophil.** *S.aureus* is within a phagosome at the start of the movie. PHAkt-eGFP recruits to the phagosome. PHAkt-eGFP diminishes from the phagosome membrane and *S.aureus* is released from the phagosome onto the surface of the neutrophil. *S.aureus* is recaptured by the neutrophil and PHAkt-eGFP re-recruits to the phagosome membrane (1^st^ pulse). Yellow cage shows the % of the bacteria which is expelled from the phagosome. Green = PHAkt-eGFP. Red = pHrodo™ red stained *S.aureus*.

**Figure S7 *S.aureus* is repeatedly released and recaptured by human neutrophils.** *S.aureus* is extracellular (in the tissue) at the start of the movie (white circle). A neutrophil phagocytoses *S.aureus* and PHAkt-eGFP recruits to the phagosome membrane (1m 36s). *S.aureus* is then released from the phagosome (5m 7s). Neutrophil then rephagocytose *S.aureus* (5m 39s) (red dot) (1^st^ pulse). *S.aureus* is then released from the phagosome (7m 3s). Neutrophil then rephagocytose *S.aureus* (7m 26s) (red dot) (2^nd^ pulse). *S.aureus* is then released from the phagosome (8m 2s). Neutrophil then rephagocytose *S.aureus* (9m 53s) (red dot) (3^rd^ pulse). The fluorescence of pHrodo™ *S.aureus* increases when *S.aureus* is expelled from the phagosome into the tissue. The fluorescence of pHrodo™ *S.aureus* decreases when *S.aureus* is recaptured. Green = CellMask™ Deep Red stained neutrophil membranes. Magenta = pHrodo™ green stained *S.aureus*.

**Figure S8 PI3Kγ inhibition prevents pulsatile recruitment of PHAkt-eGFP to neutrophil phagosomes.** *S.aureus* is within a phagosome at the start of the movie (white circle). *S.aureus* is then released from the phagosome (28s). Neutrophil then attempts to rephagocytose *S.aureus* (1m 43s) (blue dot) but PHAkt-eGFP does not re-recruit (pulse) to the phagosome. Green = PHAkt-eGFP. Magenta = pHrodo™ red stained *S.aureus*.

## STAR Methods

### RESOURCE AVAILABILITY

#### Lead contact

Further information and requests for resources and reagents should be directed to and will be fulfilled by the lead contact, Clare Muir.

#### Materials availability

*Tg(mpx:Lifeact-Ruby)sh608* was made by injecting single cell larvae with the destination plasmid, Tol2-mpx:Lifeact-Ruby (a gift from Anna Huttenlocher) with the tol2 transposase RNA generated as previously described ^37^.

#### Data and code availability

Microscopy data reported in this paper will be shared by the lead contact upon request.

All original code has been deposited in the GitHub repository https://github.com/reyesaldasoro/RingFluorescence and is publicly available as of the date of publication. DOIs are listed in the key resources table.

Any additional information required to reanalyze the data reported in this paper is available from the lead contact upon request.

### EXPERIMENTAL MODEL DETAILS

Animal work was carried out according to guidelines and legislation set out in UK law in the Animals (Scientific Procedures) Act 1986, under Project License PP7684817. Ethical approval was granted by the University of Sheffield Local Ethical Review Panel. Adult Zebrafish and larvae were maintained in the Bateson Centre aquaria at the University of Sheffield and maintained according to standard protocols ^42^. The aquarium is a continuous re-circulating closed system with a light-dark cycle of 14/10 hours respectively and a temperature of 28°C. Existing lines were London wild-type (LWT), *Nacre* and *Tg(lyz.PHAkt-eGFP)i277*. Larvae were maintained in E3 (5 mM NaCl, 0.17 mM KCl, 0.33 mM CaCl2, 0.33 mM MgSO4) plus methylene blue (Sigma-Aldrich, 50,484) at 28°C until 3dpf.

### METHOD DETAILS

#### Bacterial strain and growth conditions

*S.aureus* strain USA 300 and *M. abscessus* strain CIP104536T morphotype smooth was were used for all experiments ^39^. Overnight cultures of *S.aureus* were started from single colonies and grown in 10mL brain heart infusion medium (Sigma, 53286) overnight at 37°C, shaking at 250rpm. In the morning, cultures were diluted 1:100 and grown for 2.5hours. Cultures were pelleted at 4500g for 15mins at 4°C (Sigma, USA 3-16KL, 11240 x 13145 rotor) and resuspended in PBS (Oxoid, BR0014 G) to 1500cfu/nL. For specific experiments, *S.aureus* was killed by heating in an 80°C water bath for 20mins. *M. abscessus* cultures were grown and prepared for infection challenges (150cfu/nl) as previously described ^40^.

#### pHrodo™ staining of bacteria

200µL of PBS/bacterial suspension was then added to 1µL of either pHrodo™ Red or Green (Thermo Fisher Scientific, P36600 or P35369), incubated at 37 °C, 100 rpm for 30mins. Stained bacteria were then washed with 1mL of PBS, 1mL 30 mM Tris pH 8.5 and final wash of 1mL of PBS before resuspending in PBS for injection.

#### Zebrafish injections

Zebrafish larvae were injected using borosilicate glass needles (World Precision, Instruments, USA, TW100-4) and a pneumatic PicoPump (PV820 World Precision Instruments, USA). Borosilicate glass needles were prepared using a horizontal micropipette puller (P-1000 Sutter Flaming/Brown™) and subsequently loaded with 7μl of inoculum prepared as above using an Eppendorf ™ Microloader™ Pipette Tip. The needle was orientated into the desired position using a micromanipulator (World Precision Instruments, USA) and a stereomicroscope (Nikon microscope SMZ 745). Fine tweezers were then used to break the needle tip and the inoculum drop was then calibrated to 1nL by using a graticule (Pyser Optics) coated with mineral oil. The 18^th^-20^th^ dorsal somite of day-3 zebrafish larvae was then injected and the infection site was time-lapsed from 30 mpi.

#### Drug treatment of zebrafish

Day 3 zebrafish larvae were immersed in 1µM AS605240 and 100 µM CAL-101 (Selleckchem) in 1% DMSO in E3 medium. Larvae were incubated in inhibitors at 28°C for 30mins before imaging. Controls were similarly immersed in 1% DMSO in E3.

#### Microscopy

Larvae were mounted in a 1% low melting point agarose solution (Affymetrix, 32,830) containing 0.168 mg/mL tricaine. Agarose was covered with 500μl of clear E3 solution. Imaging was performed from 30min until 3hours post-injury using a 40x oil objective (UplanSApo 40x oil [NA 1.3]) on an UltraVIEWVoX spinning disk confocal laser imaging system (Perkin Elmer) with a Hamamatsu C9100-50 EM-CCD camera. Fluorescence for eGFP and pHrodo™ Green was acquired using an excitation wavelength of 488nm, emission was detected at 510nm. Fluorescence for pHrodo™ Red and mRuby was acquired using 525nm emission and detected at 640nm. 25-40µM z stacks were captured with a step interval of 1.5µM. Live imaging of human neutrophils were performed on the Cairnfocal system as previously described ^41^, using excitation wavelength of 470 and 647nm. Neutrophils were maintained at 37°C using using the UNO-T Stage-Top incubator (Okolab, Naples, Italy).

#### Image Analysis

Images were processed using Fiji and Matlab® (Mathworks™, Natick, USA). Phagosomes were analysed for a mean of 72mins (6.4-191.7mins +/− SD 56.7). Phagosomes were tracked using TrackMate ^43^. On average, phagosomes were analysed for 72 mins (6.4-191.7mins +/− SD 56.7). The position of the phagosomes was obtained using the tracking plug-in TrackMate ^43^ and all subsequent analysis was performed in Matlab. The volumetric analysis of the phagosomes was performed with an intensity based-segmentation on each of the 3D volumes. All channels were smoothed in 3D with a box kernel of size [3×3] and then intensity levels were used to perform an intensity-based segmentation. For the green channel two intensity levels were employed, one low that corresponded to the whole neutrophil and one high that corresponded to the phagosome. The red channel only required one level that corresponded to the bacteria. The volumes of the bacteria inside and outside the neutrophil were measured and the ratio of inside and outside was calculated as well as the average intensities for all frames. To display the volumes, isosurfaces were calculated and displayed as meshes (Fig. 3A). To calculate the intensities of the phagosomes over time, the positions were recorded as previously mentioned and then the intensities of all pixels within a ring of radius of 6 pixels were extracted. The average intensity inside the ring was calculated as well as the intensities as a function of the angle, in particular 21 directions were calculated, to visualise if the intensity was uniform around the ring or if it was higher in one side than in the other. For this study 9 phagosomes in four different datasets were analysed. Each track consisted of a 2D dataset of dimensions [number of time frames x 21] where the 21 corresponded to the angles of the ring. The intensity values were low pass filtered to smooth the results (Fig. 1A). Finally, the intensity values of the 9 tracks were averaged to calculate mean and standard deviation. First, intensities were normalised by subtracting the minimum value of the track and then dividing by the maximum so that the range of values was within 0 and 1. Next, the tracks were shifted in time so that all the start of all tracks coincided in a new time 0 and could be compared. Then mean and standard deviation per time frame were calculated (Fig.1A). The code is publicly available through GitHub in the repository: https://github.com/reyesaldasoro/RingFluorescence.

#### Preparation of human neutrophils

Human neutrophils were isolated by Plasma-Percoll density gradient centrifugation from whole blood of healthy donors as previously described ^44^. The study was carried out with written informed consent obtained from each donor. The ethical approval was obtained from the South Sheffield Research Ethics Committee (study number STH13927). Freshly isolated neutrophils were resuspended at 5×10^6^ cells/mL in phenol red-free Roswell Park Memorial Institute (RPMI) 1640 media (Thermo Fisher, Waltham, MA), supplemented with 10% (v/v) heat-inactivated fetal bovine serum (FBS, PromoCell, Heidelberg, Germany) and 25mM HEPES buffer solution (Sigma Aldrich, Gillingham, Dorset). Neutrophils were cultured on high 35mm µ-dish with ibiTreat #1.5 polymer coverslip (Ibidi, Martinsried, Germany). Freshly isolated neutrophils (5×10^5^ cells) were stained with 10 µg/mL of CellMask Deep Red plasma membrane stain (C10046, Invitrogen, Carlsbad, CA).

#### Quantification and statistical analysis

Prism software (GraphPad, Inc.) was used to perform statistical analysis. Precision measures, n values and statistical tests used are indicated in all figure legends. P = <0.05 was used to determine significance.

## Acknowledgements

We would like to acknowledge the research support teams at the University of Sheffield. We thank the aquarium staff for the breeding and maintenance of our zebrafish strains as well as the Wolfson Light Microscopy Facility. We would also like to thank Professor Anna Huttenlocher for her gift of the Tol2-mpx:Lifeact-Ruby plasmid. This work was supported by the Wellcome Trust (R/151624).

## Author contributions

Conceptualisation, C.F.M., F.E.E., A.M.C and S.A.R.; Methodology, C.F.M., F.E.E., Y.X.H., T.K.P., S.E., A.B., C.A.L., C.C.A., A.J.C., L.R.P., J.S.K., A.M.C and S.A.R.; Investigation, C.F.M., Y.X.H., K.A.B and C.C.R; Writing – Original Draft, C.F.M; Writing – Review & Editing, C.F.M., F.E.E., Y.X.H., T.K.P., A.B., C.A.L., C.C.A., A.J.C., L.R.P., J.S.K., A.M.C and S.A.R. Funding Acquisition, C.F.M., F.E.E., A.M.C and S.A.R.; Resources, J.S.K., A.M.C and S.A.R.; Supervision, A.M.C and S.A.R.

## Declaration of interests

The authors declare no competing interests.

## Notes

https://github.com/reyesaldasoro/RingFluorescence

### Competing Interest Statement

The authors have declared no competing interest.

## References

1. Welch, H.C.E., Condliffe, A.M., Milne, L.J., Ferguson, G.J., Hill, K., Webb, L.M.C., Okkenhaug, K., Coadwell, W.J., Andrews, S.R., Thelen, M., et al. (2005). P-Rex1 Regulates Neutrophil Function. Curr. Biol. 15, 1867–1873. 10.1016/J.CUB.2005.09.050.

2. Zhan, Y., Virbasius, J. V, Song, X., Pomerleau, D.P., and Zhou, G.W. (2001). The p40 phox and p47 phox PX Domains of NADPH Oxidase Target Cell Membranes via Direct and Indirect Recruitment by Phosphoinositides*. 10.1074/jbc.M109520200.

3. Schreiber, A., Rolle, S., Peripelittchenko, L., Rademann, J., Schneider, W., Luft, F.C., and Kettritz, R. (2010). Phosphoinositol 3-kinase-γ mediates antineutrophil cytoplasmic autoantibody-induced glomerulonephritis. Kidney Int. 77, 118–128. 10.1038/KI.2009.420.

4. Hoenderdos, K., Lodge, K.M., Hirst, R.A., Chen, C., Palazzo, S.G.C., Emerenciana, A., Summers, C., Angyal, A., Porter, L., Juss, J.K., et al. Hypoxia upregulates neutrophil degranulation and potential for tissue injury. 10.1136/thoraxjnl-2015-207604.

5. Schlam, D., Bagshaw, R.D., Freeman, S.A., Collins, R.F., Pawson, T., Fairn, G.D., and Grinstein, S. (2015). ARTICLE Phosphoinositide 3-kinase enables phagocytosis of large particles by terminating actin assembly through Rac/Cdc42 GTPase-activating proteins. Nat. Commun. 6. 10.1038/ncomms9623.

6. Bohdanowicz, M., Cosío, G., Backer, J.M., and Grinstein, S. (2010). Class I and class III phosphoinositide 3-kinases are required for actin polymerization that propels phagosomes. J. Cell Biol. 191, 999–1012. 10.1083/jcb.201004005.

7. Dewitt, S., Tian, W., and Hallett, M.B. (2006). Localised PtdIns(3,4,5)P3 or PtdIns(3,4)P2 at the phagocytic cup is required for both phagosome closure and Ca2+ signalling in HL60 neutrophils. J Cell Sci 119, 443–451. 10.1242/jcs.02756.

8. Marshall, J.G., Booth, J.W., Stambolic, V., Mak, T., Balla, T., Schreiber, A.D., Meyer, T., and Grinstein, S. (2001). Restricted accumulation of phosphatidylinositol 3-kinase products in a plasmalemmal subdomain during Fcγ receptor-mediated phagocytosis. J. Cell Biol. 153, 1369– 1380. 10.1083/jcb.153.7.1369.

9. Yeung, T., and Grinstein, S. (2007). Lipid signaling and the modulation of surface charge during phagocytosis. Immunol. Rev. 219, 17–36.

10. Wang, X., Robertson, A.L., Li, J., Chai, R.J., Haishan, W., Sadiku, P., Ogryzko, N. V., Everett, M., Yoganathan, K., Luo, H.R., et al. (2014). Inhibitors of neutrophil recruitment identified using transgenic zebrafish to screen a natural product library. Dis. Model. Mech. 7, 163–169. 10.1242/dmm.012047.

11. Greene, C.J., Nguyen, J.A., Cheung, S.M., Arnold, C.R., Balce, D.R., Wang, Y.T., Soderholm, A., McKenna, N., Aggarwal, D., Campden, R.I., et al. (2022). Macrophages disseminate pathogen associated molecular patterns through the direct extracellular release of the soluble content of their phagolysosomes. Nat. Commun. 13, 1–17. 10.1038/s41467-022-30654-4.

12. Wang, X., Robertson, A.L., Li, J., Chai, R.J., Haishan, W., Sadiku, P., Ogryzko, N.V., Everett, M., Yoganathan, K., Luo, H.R. and Renshaw, S.A. (2014). Inhibitors of neutrophil recruitment identified using transgenic zebrafish to screen a natural product library. Co. Biol. 7, 163–169. 10.1242/dmm.012047.

13. Pazhakh, V.I., Ellett, F., Croker ID, B.A., O, J.A., Pase, L., SchulzeID, K.E., Stefan Greulich, R., Gupta, A., Carlos Reyes-AldasoroID, C., Andrianopoulos, A., et al. (2019). β-glucan-dependent shuttling of conidia from neutrophils to macrophages occurs during fungal infection establishment. 10.1371/journal.pbio.3000113.

14. Hellebrekers, P., Hietbrink, F., Vrisekoop, N., Leenen, L.P.H., and Koenderman, L. (2017). Neutrophil functional heterogeneity: Identification of competitive phagocytosis. Front. Immunol. 8, 1–9. 10.3389/fimmu.2017.01498.

15. Hawkins, P.T., and Stephens, L.R. (2014). PI3K signalling in inflammation ☆. BBA - Mol. Cell Biol. Lipids 1851, 882–897. 10.1016/j.bbalip.2014.12.006.

16. Yoo, S.K., Deng, Q., Cavnar, P.J., Wu, Y.I., Hahn, K.M., and Huttenlocher, A. (2010). Differential Regulation of Protrusion and Polarity by PI(3)K during Neutrophil Motility in Live Zebrafish. Dev. Cell 18, 226–236. 10.1016/j.devcel.2009.11.015.

17. Sadhu, C., Masinovsky, B., Dick, K., Sowell, C.G., and Staunton, D.E. (2003). Essential role of phosphoinositide 3-kinase delta in neutrophil directional movement. J. Immunol. 170, 2647–2654. 10.4049/jimmunol.170.5.2647.

18. Sadhu, C., Dick, K., Tino, W.T., and Staunton, D.E. (2003). Selective role of PI3Kδ in neutrophil inflammatory responses. Biochem. Biophys. Res. Commun. 10.1016/S0006-291X(03)01480-3.

19. Elworthy, S., Rutherford, H.A., Prajsnar, T.K., Hamilton, N.M., Vogt, K., Renshaw, S.A., and Condliffe, A.M. (2023). Activated PI3K delta syndrome 1 mutations cause neutrophilia in zebrafish larvae. DMM Dis. Model. Mech. 16. 10.1242/dmm.049841.

20. Gilbert, A.S., Seoane, P.I., Sephton-Clark, P., Bojarczuk, A., Hotham, R., Giurisato, E., Sarhan, A.R., Hillen, A., Velde, G. Vande, Gray, N.S., et al. (2017). Vomocytosis of live pathogens from macrophages is regulated by the atypical MAP kinase ERK5. Sci. Adv. 3, e1700898. 10.1126/sciadv.1700898.

21. Johnston, S.A., and May, R.C. (2010). The human fungal pathogen Cryptococcus neoformans escapes macrophages by a phagosome emptying mechanism that is inhibited by arp2/3 complex-mediated actin polymerisation. PLoS Pathog. 6, 27–28. 10.1371/journal.ppat.1001041.

22. Bain, J.M., Lewis, L.E., Okai, B., Quinn, J., Gow, N.A.R., and Erwig, L.P. (2012). Non-lytic expulsion/exocytosis of Candida albicans from macrophages. Fungal Genet. Biol. 49, 677– 678. 10.1016/j.fgb.2012.01.008.

23. Alvarez, M., and Casadevall, A. (2006). Phagosome Extrusion and Host-Cell Survival after Cryptococcus neoformans Phagocytosis by Macrophages. Curr. Biol. 16, 2161–2165. 10.1016/j.cub.2006.09.061.

24. Ma, H., Croudace, J.E., Lammas, D.A., and May, R.C. (2006). Expulsion of Live Pathogenic Yeast by Macrophages. Curr. Biol. 16, 2156–2160. 10.1016/j.cub.2006.09.032.

25. Fu, M.S., Coelho, C., De Leon-Rodriguez, C.M., Rossi, D.C.P., Camacho, E., Jung, E.H., Kulkarni, M., and Casadevall, A. (2018). Cryptococcus neoformans urease affects the outcome of intracellular pathogenesis by modulating phagolysosomal pH 10.1371/journal.ppat.1007144.

26. Smith, L.M., Dixon, E.F., and May, R.C. (2015). The fungal pathogen Cryptococcus neoformans manipulates macrophage phagosome maturation. Cell. Microbiol. 17, 702–713. 10.1111/cmi.12394.

27. P Cech, R.L. (1984). Heterogeneity of human neutrophil phagolysosomes: functional consequences for candidacidal activity. Blood 64, 147–151.

28. Servant, G., Weiner, O.D., Herzmark, P., Balla, T., Sedat, J.W., and Bourne, H.R. (2000). Polarization of Chemoattractant Receptor Signaling During Neutrophil Chemotaxis. Science (80-.). 287, 1037–1040.

29. Quinn, S.E., Huang, L., Kerkvliet, J.G., Swanson, J.A., Smith, S., Hoppe, A.D., Anderson, R.B., Thiex, N.W., and Scott, B.L. (2021). The structural dynamics of macropinosome formation and PI3-kinase-mediated sealing revealed by lattice light sheet microscopy. Nat. Commun. 2021 121 12, 1–12. 10.1038/s41467-021-25187-1.

30. Suire, S., Condliffe, A.M., Ferguson, G.J., Ellson, C.D., Guillou, H., Davidson, K., Welch, H., Coadwell, J., Turner, M., Chilvers, E.R., et al. (2006). Gβγs and the Ras binding domain of p110γ are both important regulators of PI3Kγ signalling in neutrophils. Nat. Cell Biol. 8, 1303–1309. 10.1038/ncb1494.

31. Deladeriere, A., Gambardella, L., Pan, D., Anderson, K.E., Hawkins, P.T., and Stephens, L.R. (2015). The regulatory subunits of PI3Kγ control distinct neutrophil responses. Sci. Signal. 8. 10.1126/scisignal.2005564.

32. Rathinaswamy, M.K., Fleming, K.D., Dalwadi, U., Pardon, E., Harris, N.J., Yip, C.K., Steyaert, J., and Burke, J.E. (2021). HDX-MS-optimized approach to characterize nanobodies as tools for biochemical and structural studies of class IB phosphoinositide 3-kinases. Structure 29, 1371–1381.e6. 10.1016/j.str.2021.07.002.

33. Cox, D., Tseng, C.C., Bjekic, G., and Greenberg, S. (1999). A requirement for phosphatidylinositol 3-kinase in pseudopod extension. J. Biol. Chem. 274, 1240–1247. 10.1074/jbc.274.3.1240.

34. Hirsch, E., Katanaev, V.L., Garlanda, C., Azzolino, O., Pirola, L., Silengo, L., Sozzani, S., Mantovani, A., Altruda, F., and Wymann, M.P. (2000). Central role for G protein-coupled phosphoinositide 3-kinase γ in inflammation. Science (80-.). 287, 1049–1052. 10.1126/SCIENCE.287.5455.1049/SUPPL_FILE/1044275S1_THUMB.GIF.

35. Vieira, O. V., Botelho, R.J., Rameh, L., Brachmann, S.M., Matsuo, T., Davidson, H.W., Schreiber, A., Backer, J.M., Cantley, L.C., and Grinstein, S. (2001). Distinct roles of class I and class III phosphatidylinositol 3-kinases in phagosome formation and maturation. J. Cell Biol. 155, 19–25. 10.1083/jcb.200107069.

36. Dbouk, H.A., Vadas, O., Shymanets, A., Burke, J.E., Salamon, R.S., Khalil, B.D., Barrett, M.O., Waldo, G.L., Surve, C., Hsueh, C., et al. (2012). G protein-coupled receptor-mediated activation of p110b by Gβγ is required for cellular transformation and invasiveness. Sci. Signal. 5. 10.1126/scisignal.2003264.

37. Kwan, K.M., Fujimoto, E., Grabher, C., Mangum, B.D., Hardy, M.E., Campbell, D.S., Parant, J.M., Yost, H.J., Kanki, J.P., and Chien, C. Bin (2007). The Tol2kit: A multisite gateway-based construction Kit for Tol2 transposon transgenesis constructs. Dev. Dyn. 236, 3088– 3099. 10.1002/dvdy.21343.

38. Lam, S.H., Chua, H.L., Gong, Z., Lam, T.J., and Sin, Y.M. (2004). Development and maturation of the immune system in zebrafish, Danio rerio: a gene expression profiling, in situ hybridization and immunological study. Dev. Comp. Immunol. 28, 9–28.

39. Fey, P.D., Endres, J.L., Yajjala, V.K., Widhelm, T.J., Boissy, R.J., Bose, J.L., and Bayles, K.W. (2013). A Genetic Resource for Rapid and Comprehensive Phenotype Screening of Nonessential Staphylococcus aureus Genes. MBio 4. 10.1128/mBio.00537-12.

40. Bernut, A., Dupont, C., Sahuquet, A., Herrmann, J.L., Lutfalla, G., and Kremer, L. (2015). Deciphering and Imaging Pathogenesis and Cording of Mycobacterium abscessus in Zebrafish Embryos. J. Vis. Exp. 2015, 53130. 10.3791/53130.

41. Ho, Y.X., Steele, E., Prince, L., and Cadby, A. (2021). Multi-modal imaging reveals dynamic interactions of Staphylococcus aureus within human neutrophils. bioRxiv, 2021.03.05.434126.

42. Aleström, P., D’Angelo, L., Midtlyng, P.J., Schorderet, D.F., Schulte-Merker, S., Sohm, F., and Warner, S. (2020). Zebrafish: Housing and husbandry recommendations. Lab. Anim. 54, 213–224. 10.1177/0023677219869037.

43. Tinevez, J.Y., Perry, N., Schindelin, J., Hoopes, G.M., Reynolds, G.D., Laplantine, E., Bednarek, S.Y., Shorte, S.L., and Eliceiri, K.W. (2017). TrackMate: An open and extensible platform for single-particle tracking. Methods 115, 80–90. 10.1016/j.ymeth.2016.09.016.

44. Prince, L.R., Allen, L., Jones, E.C., Hellewell, P.G., Dower, S.K., Whyte, M.K.B., and Sabroe, I. (2004). The role of interleukin-1β in direct and toll-like receptor 4-mediated neutrophil activation and survival. Am. J. Pathol. 165, 1819–1826. 10.1016/S0002-9440(10)63437-2.

